# Role of tonsillar chronic inflammation and commensal bacteria in the pathogenesis of pediatric OSA

**DOI:** 10.1101/2021.01.26.428230

**Authors:** Lindybeth Sarmiento Varón, Javier De Rosa, Raquel Rodriguez, Pablo M. Fernández, L. Ariel Billordo, Plácida Baz, Gladys Beccaglia, Nicolás Spada, F. Tatiana Mendoza, Claudia M. Barberis, Carlos Vay, M. Elena Arabolaza, Bibiana Paoli, Eloísa I. Arana

## Abstract

Immune responses at the boundary between the host and the world beyond are complex and mucosal tissue homeostasis relies on them. Obstructive sleep apnea (OSA) is a syndrome suffered by children with hypertrophied tonsils. We uncovered a crucial role of pro-inflammatory tonsillar B and T cells in sustaining hypertrophy and hyperplasia by producing TNF and IL17, respectively. We detected prominent levels of expression of CD1d by tonsillar stratified as well as reticular epithelium, which have not previously been reported. By combining bacterial culture from the tonsillar core and subsequent identification of the respective isolates, we determined the most prevalent species within the cohort of OSA patients. Although the isolated species are considered normal oropharyngeal commensals in children, we confirmed their capacity to breach the epithelial barrier. Our work shed light on the pathological mechanism underlying OSA, highlighting the relevance taken by the host immune system when defining infection versus colonization.

## Introduction

Human paired palatine tonsils are lymphoid epithelial tissues of the oral mucosa around the oropharynx. They are part of the Waldeyer’s ring of lymphoid tissue, which also comprise the adenoids and the lingual tonsils. The palatine tonsils (tonsils, from now on) are strategically located to generate mucosal immunity as they are constantly exposed to dietary and airborne antigens (Ags). Moreover, they have evolved for direct transport of foreign material from the exterior to the lymphoid cells through deep and branched crypts and the absence of Ag-degrading digestive enzymes (Brandtzaeg, 2013). The surfaces of the human body, including the oropharynx, are colonized by several microbes, mainly bacteria, that establish a mutualistic relationship with the host. Tonsils are predominantly B-cell organs, immunologically most active between 4 and 10 years old. Some children (also some adults, but were not included in this study) present tonsillar hyperplasia and hypertrophy for yet unknown reasons. Such enlargement is the major pathophysiological sign underlying OSA, a highly prevalent disease, recognized as a major public health burden. OSA is characterized by repeated events of partial or complete upper airway obstruction during sleep that lead to disruption of normal ventilation with all the consequences implied due to hypoxemia. An increment in pro-inflammatory cytokines have been reported in blood from OSA patients (Ye *et al*, 2012), (Huang *et al*, 2016), (Kheirandish-Gozal & Gozal, 2019). It is assumed that the B cell hyperplasia and hypertrophy that cause OSA are the result of chronic inflammation of palatine tonsils in children.

B cells contribute to immune responses during infectious, inflammatory and autoimmune diseases. In the last two decades, we have learned that B cells are able to modulate physiological and pathological processes not only by producing antibodies (Abs) and presenting Ags but also by producing cytokines. It has been shown that B cells can occur in the form of several cytokine-secreting subsets with either pro- or anti-inflammatory functions (Fillatreau, 2018). Being an important reservoir of human B cells, the tonsils serve as a platform to study such subsets. Within this framework, we have recently demonstrated that OSA tonsils rendered significantly lower percentages of interleukin 10 (IL10) producing B cells (Bregs) than tonsils excised due to recurrent tonsillitis, showing that Bregs have a more complex and interesting role in tonsillar disease than was hitherto appreciated. Moreover, such defect in Breg population correlated with an increment in the proportion of germinal center (GC) cells at which expense tonsils are hypertrophied, revealing a role for the Breg subset in controlling GC reactions (Sarmiento Varon et al., 2017). GCs are the microanatomical sites within secondary lymphoid organs, critical for memory B and plasma cell generation. There are also many other cellular and molecular players involved in controlling GC activity. For instance, it has been long known that the pro-inflammatory cytokine tumor necrosis factor alpha (TNFa) is a required autocrine B-cell growth factor (Boussiotis *et al*, 1994).

In the present paper, we found that tonsillar mononuclear cells (TMC) from OSA tonsils effectively exhibit a pro-inflammatory cytokine profile rapidly in culture. Interestingly, under certain stimulating conditions, B lymphocytes became the main cell population driving TNFa levels in culture *ex vivo.* On the other hand, tonsillar interleukin 17 (IL17) was produced primarily by CD4^+^ T cells. At tissue level, we discovered CD1d expression by tonsillar stratified as well as reticular epithelium and corroborated the persistent hypertrophy of GC and the concomitant hyperplasia of B lymphocytes, prevalently IgG/IgM positive (IgG/IgM)^+^. We identified a number of bacterial species in the tonsillar core of patients with tonsillar hypertrophy, considered normal oropharyngeal commensals in children. However, we confirmed their presence beyond the epithelial boundaries by fluorescence *in situ* hybridization (FISH). Thus, our observations suggest that tonsillar hypertrophy is a multifaceted condition not associated to the presence of a particular microorganism but more likely to a failure of normal immune homeostatic mechanisms caused by the local loss of the capacity to discriminate between commensals and pathogens of the host. All in all, when such discrimination is lost, commensals become pathogens. Therefore, we support the notion that OSA in children is of infectious nature, clearly not associated to a single species.

## Results

### Characterization of TNFα production by B cells from hypertrophied tonsils

It is long established that B-cell–derived TNFa plays a crucial role in the development of follicular dendritic cells (FDCs) and B-cell follicles in spleen (Fu & Chaplin, 1999);(Endres *et al*, 1999)’(Gonzalez *et al*, 1998). TNFa and IL10 are cytokines with antagonistic actions. Interestingly, tonsils from OSA patients present a defective Breg compartment which correlates with higher proportion of GC B lymphocytes (BGC (Sarmiento Varon *et al*, 2017)), larger GC (Reis *et al*, 2013), and increased numbers of T follicular helper cells (Tfh) (Yamashita *et al*, 2016) than tonsils excised by other pathologies. Taking all into account, it would be expected that OSA tonsils exhibit a significant lymphocyte–derived TNFa compartment. We examined TNFa expression at the single cell level, by fluorescence-activated cell sorting (FACS), upon TMC culture. A set of cultures was treated for 24 hs with IL2 and IL4 (IL2+IL4), which promote survival of all lymphocyte subsets through slight stimulation (Perez *et al*, 2014). Another set of cultures was supplemented for 24 hs and 48 hs with CpG and CD40L (in addition to IL2 and IL4, IL2+IL4+CD40L+CpG) to target stimulation towards B cells, aiming to assess specifically their capacity to contribute to the tonsillar pool of TNFa. Single cells were selected based on FSC-A vs FSC-H analysis (Singlets gate, Fig. 1A). The cells that were already dead prior to permeabilization were excluded from the analysis based on eFluor 780 dead cell staining (Viable gate, Fig. 1A). In agreement with previous observations (Perez *et al.,* 2014), elevated levels of cell death were linked to stimulation of tonsillar cultures (Fig. 1A and B), dominated by a variety of terminally differentiated and highly activated B cells. In fact, we used this decay in the viability of the TMC cultures to monitor for suitable activation. In our experience, TMC culture stimulation is quite susceptible to minimal changes in experimental conditions (cellular density or FCS and activation cocktail batches, for instance). Therefore, when comparing cytokine secretion within a cohort of patients, those cultures that did not present such evolution in terms of viability, proportion of CD3^+^ and CD20^+^ populations and CD20 downmodulation (Fig. 1A), were not considered. Also, CD3^+^ cells served as an internal control for cytokine expression (Fig. 1C). As expected, we observed that TMC from hypertrophied tonsils produced considerable amounts of TNFa when stimulated. At 24 hs post stimulation, approximately one third (29% ± SD 22%) of the cells from the IL2+IL4 stimulated cultures, expressed TNFα and that percentage reached ~40% (37%± SD 16%) in CD40L+CpG+IL2+IL4 stimulated cultures. The latter cultures were particularly affected by the surface CD20 down-modulation which takes place in response to general stimuli (Valentine *et al*, 1987), albeit more pronounced with CD40L stimulation, as it has been extensively described previously (Anolik *et al*, 2003)’(Sarmiento Varon *et al*, 2017). Interestingly, B cells (CD20^+^) expressing TNFa were those presenting lower levels of CD20 (CD20^down^ TNFa ^+^ cells, Fig. 1A) suggesting that TNFa expression might be another functional consequence of CD20 modulation and the downstream signalling pathways triggered post internalization. These findings confirmed that the extent of CD20 down-regulation correlates with the degree of B cell activation (Fig. 1A and C) (Valentine *et al.,* 1987)’(Anolik *et al.,* 2003)’(Sarmiento Varon *et al*, 2017).

**Figure 1.**
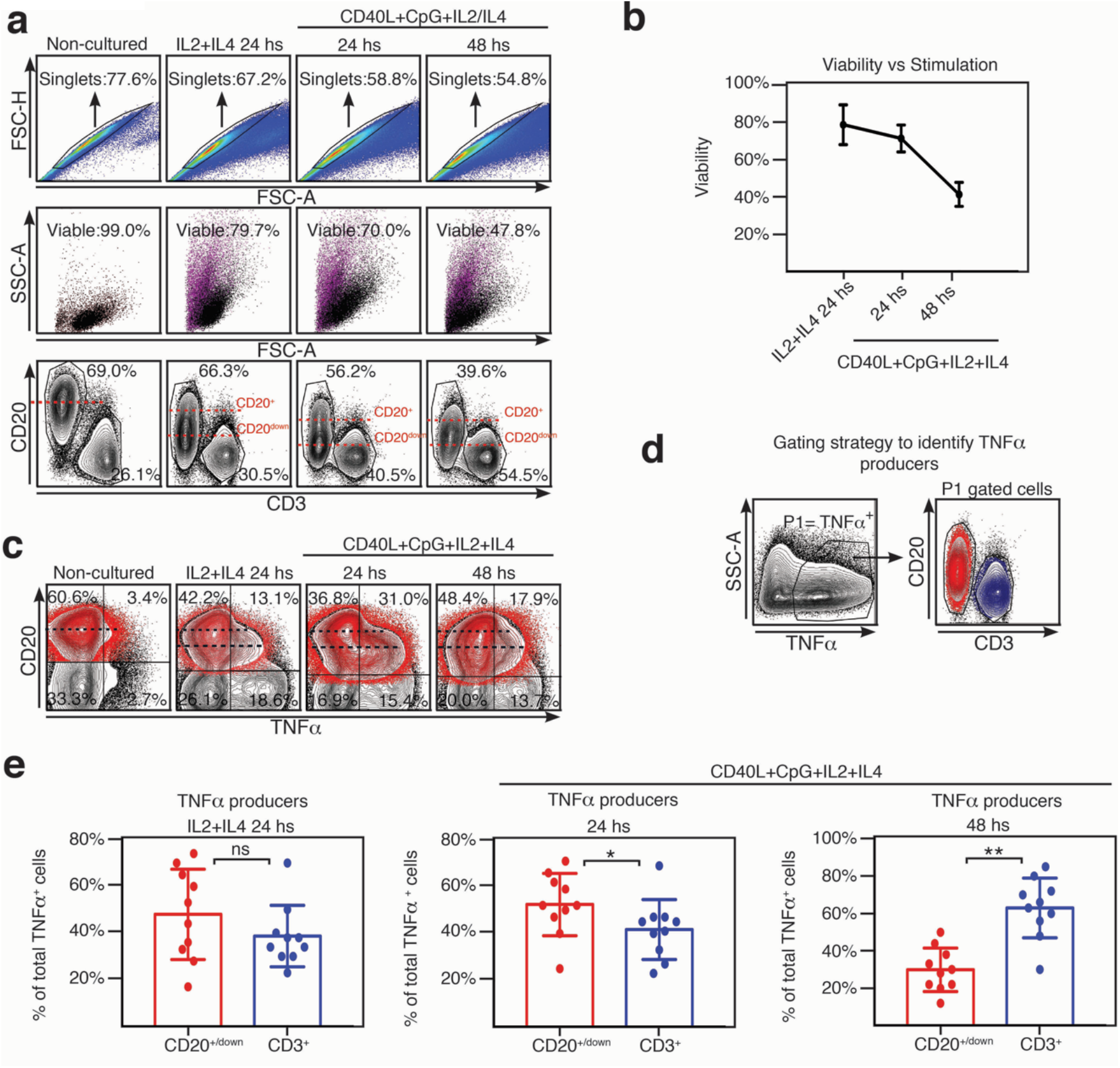
TNF expression by tonsillar mononuclear cells. A) Freshly isolated TMC were cultured xon IL 2+IL 4 alone or CpG+CD40L+IL 2+IL 4, for the time points indicated on the top of each panel either. Non-cultured cells served as control (conserved at 4^0^C). Samples were subsequently analyzed by FACS. Gating strategy is illustrated. Singlets were gated by plotting FSC-A vs FSC-H for each sample (upper panels). Within the singlets population, purple color denotes dead cells determined by eFluor 780 staining. The viable gate corresponds to the eFluor 780^-^cells (depicted in black, middle panels). To detect TNFa in cultured cells, they were stimulated with PMA/ionomycin/Brefeldin A for the last 5 hs. They were then stained for surface CD20, CD3 and intracellular TNFa. Percentages designate frequencies of the populations indicated. Dashed red lines show the decline of expression of CD20 post stimulation. Data from one experiment representative of 10 independent experiments performed with different individuals each one, are presented. B) Line graph plotting the mean percentage ± SD of the frequencies of the viable gate pooled from the 10 independent experiments performed with stimulated TMC and detailed in A). C) Freshly isolated TMC were cultured on IL 2+IL 4 alone or CpG+CD40L+IL 2+IL 4, for the time points indicated on the top of each panel. Non-cultured cells served as control. To detect TNFa, cells from all the panels were stimulated with PMA/ionomycin/Brefeldin A for the last 5 hs (included non-cultured), subsequently stained for surface CD20, CD3 and intracellular TNFa and finally analyzed by FACS. Contour plots show TNFa and CD20 expression by TMC. They correspond to events within singlets and viable gates illustrated in A). Percentages designate frequencies of populations in each quadrant. CD20^+/down^ cells appear highlighted in red, based on the gating performed on the corresponding CD20 vs CD3 graph as in A), using the backgating option, to exclude any CD3 contamination. Dashed black lines show the decline of expression of CD20 post stimulation. Data from one experiment representative of 10 independent experiments performed with different individuals each one, are presented. D) Gating strategy used to estimate B and T cell specific contribution to the TNFa^+^ cell pool from the 10 samples analyzed as shown in A), B), C). P1 denotes percentage of TNFa^+^ cell population determined by TNFa vs SSC-A contour profile under the stimulations described above. Panel on the right, representative contour plots for CD20^+/down^ (red) and CD3^+^ (blue) populations within the P1 gate. E) Histograms presenting the mean percentage ± SD of the CD3^+^ and CD20^down/+^ cell population frequencies from 10 independent experiments within the TNFa^+^ cell pool for each stimulating condition, calculated as shown in D) *\*p* < 0.05 and \*\**p* < 0.01, unpaired *t* test.

We next determined in which proportion CD20^+^ and CD3^+^ cells were contributing to the total TNF a^+^ population (Fig. 1D). Comprehensibly, at 24 hs post stimulation with CD40L+CpG+IL2+IL4, when B cell activation peaked, CD20^+/down^ cells represented the majority of tonsillar TNFa ^+^ cells (52.4% ± SD 13.4% CD20^+/down^ cells vs 41.7% ± SD 12.8% CD3^+^ cells). On the other hand, at 48 hs post stimulation with CD40L+CpG+IL2+IL4, when tonsillar B cells showed strongly reduced viability, tonsillar TNFa^+^ population was dominated by CD3^+^ cells (30.0% ± SD 11.7% CD20^+/down^ cells vs 63.0% ± SD 15.9% CD3^+^ cells). The cultures stimulated for 24 hs only with IL2+IL4 did not rendered significant differences between CD20^+/down^ and CD3^+^ cells contribution to the TNFa^+^ cell pool (Fig. 1E). Finally, we also tried stimulating the TMC cultures using as a surrogate antigen F(ab)2 goat antibodies specific for IgM and IgG (in addition to CD40L+IL2+IL4) (Perez *et al.,* 2014), obtaining very similar results to those of CD40L+CpG+IL2+IL4 stimulation (data not shown). Parekh VV *et al.* (Parekh *et al*, 2009) have previously reported that upon culture, TMC from hypertrophied tonsils rendered higher levels of TNFa^+^ in the supernatant than TMC from children with recurrent tonsillitis. These results extend such findings, indicating that being exposed to various stimuli, B cells can be the main producers of TNFa, supporting (if not promoting) tonsillar inflammation.

### Presence of inflammatory T cells co-expressing IL17 and TNFa in hypertrophied tonsils

It has been suggested that development of OSA is associated with peripheral Th17/Treg imbalance and is characterized by a pro-inflammatory cytokine microenvironment (Ye *et al.,* 2012). We have previously reported on the ability of tonsillar B cells to secrete IL17 (Sarmiento Varon *et al*, 2017). However, while B cells can be the main producers of tonsillar TNFa as we showed above, we found IL17 secretion is indeed dominated by CD3^+^ cells at every culture condition tested, including those specifically stimulating B lymphocytes (Fig. 2A and B). TMC cultures supplemented with IL2+IL4 as well as those supplemented with CD40L+CpG+IL2+IL4 rendered an IL17^+^ population which comprised ~90% CD3^+^ CD4^+^IL17^+^ cells at 24 and 48 hs. We confirmed this observation by measuring cytokines on the supernatant of some of the cultures by multiplex biomarker immunoassay (Luminex). We tried 4 different conditions to specifically stimulate B cells (CpG; CD40L; anti-(IgM/IgG)+CpG; anti-(IgM/IgG)+ CD40L), all of them in addition to IL2+IL4. We also added a culture treated only with IL2+IL4, as control. IL17 concentration did not significantly varied with culture condition compared to the control, as opposed to that of TNFa, which exhibited significant increments under B cell stimulating conditions (data not shown). As expected, CD3^+^IL17^+^ cells were confirmed to be CD3^+^CD4^+^IL17^+^ (Th17) cells (Fig. 2B).

**Figure 2.**
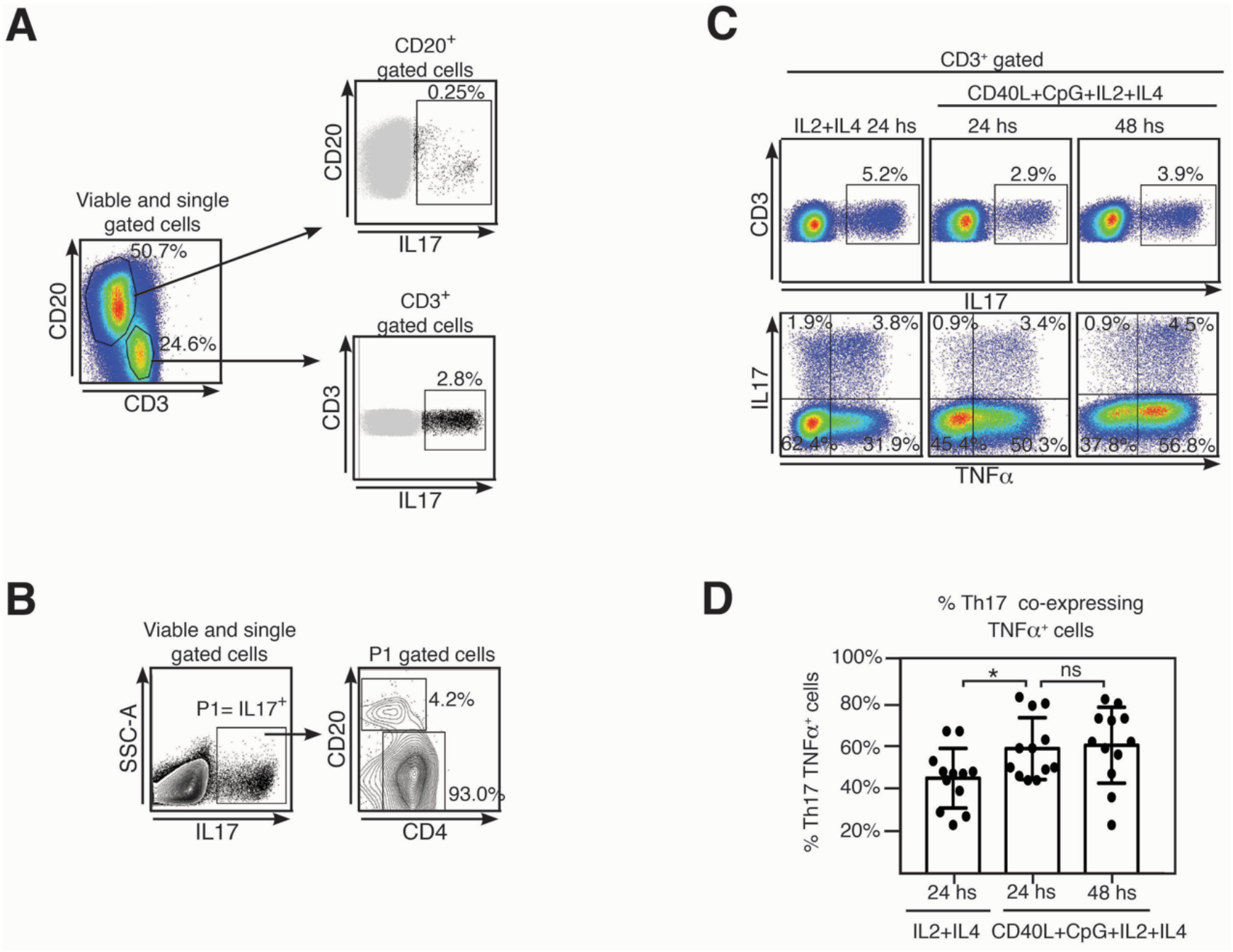
Tonsillar Th17 cells co-express TNFa and IFNγ. A) Freshly isolated TMC from a were cultured on CpG+CD40L+IL 2+IL 4 for 24 hs. Samples were subsequently analyzed by FACS. Gating strategy is illustrated. Singlets were gated by plotting FSC-A vs FSC-H for each sample. Within the singlets population, dead cells determined by eFluor 780 staining. To detect IL17 in cultured cells, they were stimulated with PMA/ionomycin/Brefeldin A for the last 5 hs. They were then stained for surface CD20, CD3 and intracellular IL17. Pseudocolor plots show CD3 and CD20 expression by TMC upon stimulation, percentages designate frequencies of populations in each gate (left panel). Upper right panel: dot plot shows IL17 and CD20 expression by TMC, CD20^+^IL17^+^ population is highlighted in black and its frequency within CD20^+^ population is indicated as percentage. Lower right panel: dot plot shows IL17 and CD3 expression by TMC, CD3^+^IL17^+^ population is highlighted in black and its frequency within CD3^+^ population is indicated as percentage. Data from one experiment representative of 10 independent experiments performed with different individuals each one, are presented. B) Gating strategy used to estimate the populations contributing to the IL17^+^ cell subset. P1 denotes L17^+^ cell population determined by TNFa vs SSC-A contour profile. Panel on the right, representative contour plots for CD20^+^ and CD4^+^ populations within the P1 gate. Percentages designate frequencies of the populations indicated. Data from one experiment representative of 6 independent experiments performed with different individuals each one, are presented C) Freshly isolated TMC were cultured on IL 2+IL 4 alone or CpG+CD40L+IL 2+IL4, for the time points indicated on the top of each panel. To detect TNFa and IL17, cells from all the panels were stimulated with PMA/ionomycin/Brefeldin A for the last 5 hs, subsequently stained for surface CD20, CD3, intracellular TNFa and IL17 and finally analyzed by FACS. Singlets were gated by plotting FSC-A vs FSC-H for each sample. Within the singlets population, dead cells determined by eFluor 780 staining. Upper panels: pseudocolour plots show IL17 vs CD3 expression by TMC. They correspond to events within singlets, viable and CD3^+^ gates. Percentages designate frequencies of IL17^+^ cells within CD3^+^. Lower panels: pseudocolour plots show TNFa and IL17 expression by TMC, gated as in the upper panels. Percentages designate frequencies of populations in each quadrant. Data from one experiment representative of 12 independent experiments performed with different individuals each one, are presented D) Histograms presenting the mean percentage ± SD of the Th17 TNFa^+^ cell population frequencies from 12 independent experiments for each stimulating condition, calculated as shown in C), top right quadrant *\*p* < 0.05 unpaired *t* test.

Th17 cells have appeared to exhibit plasticity of function and often co-produce other pro-inflammatory cytokines, particularly at sites of inflammation. Hence, we investigated whether tonsillar Th17 also expressed other cytokines as TNFa and interferon γ (IFNγ). The pseudocolor dot plots shown in Fig. 2C were gated on CD3^+^ cells and show CD3 vs IL17 (upper panel) and IL17 vs TNFa (lower panel) staining. We detected co-expression of TNFa in a significant fraction of the Th17 cell population upon all culture conditions tested. At 24 hs post stimulation, nearly half of the Th17 population (44% ± SD 14%) from the IL2+IL4 stimulated cultures co-expressed TNFa. This percentage significantly increased when TMC were cultured with CD40L+CpG+IL2+IL4 for 24 hs (59% ± SD 14,5%) as well as for 48 hs (60%± SD 18%) indicating a positive correlation between higher frequencies of TNFa and IL17 and TNFa co-expression (Fig. 2D). Low levels of expression of IFNγ observed upon the stimulating conditions tested precluded us from definitive conclusions regarding co-expression of IFNγ and IL17.

We concluded that tonsillar IL17 was produced primarily by CD4^+^ T cells, which largely coproduced TNFa. Interestingly, several studies have linked the Th17 pathway with formation of GCs in mice spleens and ectopic B cell follicles at sites of inflammation (Ding *et al*, 2013), (Barbosa *et al*, 2011) (Quinn *et al*, 2018). On the whole, TMC from hypertrophied tonsils promptly exhibited a pro-inflammatory cytokine profile in culture.

### B cell driven alterations on the complex histological milieu of tonsils

Like all mucosae, tonsillar immune actions (anti- and pro-inflammatory) must be tightly regulated, to balance the protection against virulent germs and the tolerance to harmless flora and innocuous Ags entering with air and food. Collectively, the results described above and others we have previously published (Sarmiento Varon *et al*, 2017) (Perez *et al.,* 2014) are in agreement with the notion that tonsillar hypertrophy is a result of an imbalance between regulatory and effector immune functions which in turn lead to a local chronic inflammation. To this point, we have examined isolated cells. However, *in vivo,* immune processes occur in a complex tissue context which has to be comprehended. In order to recognize the histologic compartments within tonsils, we performed immune-fluorescence staining on cryosections from excised tonsils. The palatine tonsils are secondary lymphoid organs belonging to the mucosal associated lymphoid tissue (MALT), with unique histological characteristics. Unlike the lymph nodes, tonsils do not have afferent lymphatics, but present deep and branched crypts. Ag sample occurs through their distinctive tissue architecture and this is why tonsils are called lympho-epithelial organs. The tonsillar epithelium provides protection as well as serving to transport foreign material from the lumen to the lymphoid compartment. To evidence this mucosal surface, we tested the tonsillar epithelial cells’ reactivity to antibodies against CD1d, since it has been reported its expression in other mucosal epithelial cells, like intestinal (Blumberg *et al*, 1991), epidermal keratinocytes (Bonish *et al*, 2000) and vaginal epithelium (Kawana *et al*, 2008). We confirmed CD1d expression in tonsillar epithelium. Anatomically, the surface epithelium of the palatine tonsils is an extension of the stratified squamous epithelium of the oral mucosa (Fig. 3A). Importantly, in the crypts, the stratified epithelium becomes narrower and laxly textured (reticulated epithelium as termed by Oláh in 1978 (Olah & Glick, 1978)) and highly infiltrated with lymphocytes (Fig. 3B, C and D). We observed CD1d expression in reticulated epithelium as well as in stratified squamous epithelium. Nevertheless, the level of that expression did not appear uniform at all levels of the multilayered squamous epithelium. The basal cells, which maintained an orientation perpendicular to the basement membrane, presented negligible immunoreactivity to anti CD1d. Conversely, CD1d was expressed strongly in the epithelial cells of the supra-basal and intermediate layers, significantly declining its expression in the apical layers. Expression was restrained mostly to the cell membrane in all cases (Fig. 3). This epithelium was supported by a layer of connective tissue (Fig. 3A) that separates it from the lymphoid component, and resulted negative for CD1d, with the exception of sporadic infiltrating lymphoid cells. Fig. 3B and C show a region of crypts lined by reticulated epithelium. In fact, the crypt in Fig. 3C, is lined by reticulated epithelium on one side and stratified squamous epithelium on the other. Two characteristics of the former are clear from the figures, and these are the desquamation of the upper cell layers and the absence or disruption of the basal layer of the epithelium. In this case, there is no boundary between the epithelium and the underlying lymphoid tissue. The cells in the intermediate layer were distorted and often separated from one another by the invading lymphoid cells, yet again CD1d exhibited high expression in this layer. The reticular epithelium trailed the curving shapes of the underlying follicles. To our knowledge, this is the first report on CD1d expression by tonsillar epithelium.

**Figure 3.**
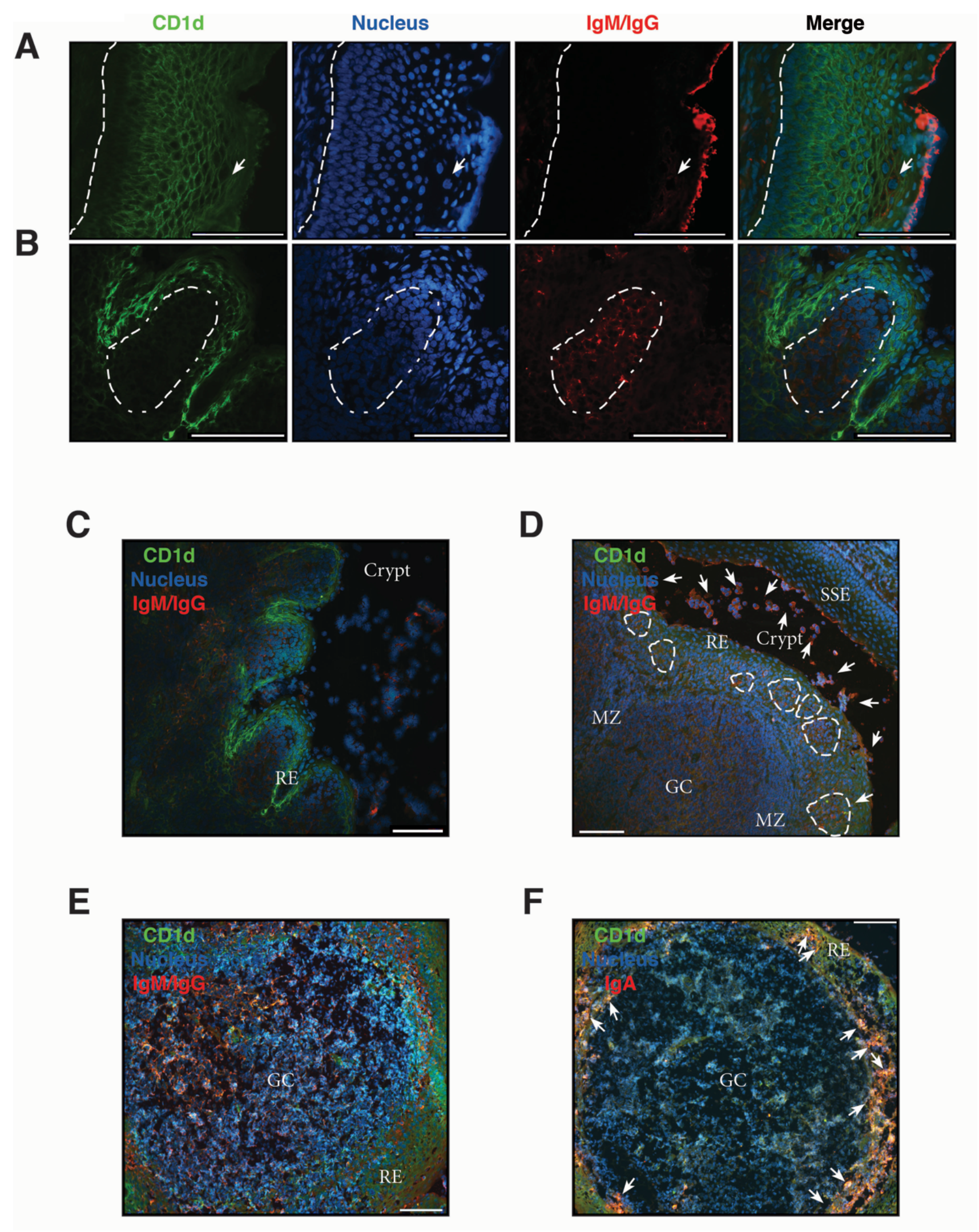
Analysis of chronically inflamed human tonsils by immunohistology. A)-F) Three-colour immunofluorescent microscopy of tonsillar frozen sections. A) Representative immunofluorescence staining showing stratified squamous epithelium with CD1d (green), IgG/IgM (red) and nucleus (blue). Dashed line represents the limit between the basal membrane and the epithelium. Arrow indicates IgG^+^/IgM^+^ plasma and B cells intercalated in the mesh of epithelial cells. B) Representative immunofluorescence staining showing reticular epithelium with CD1d (green), IgG/IgM (red) and nucleus (blue). Dashed lines represent the limit of the follicle. A) and B) Scale bar, 104 μm. C) A less magnified image of the section shown in B) to evidence the particular characteristics of the lympho-epithelium covering the crypts. RE stands for reticular epithelium. Scale bar, 104 μm. D) Representative immunofluorescence staining showing a germinal center (GC) with its mantle zone (MZ), the reticular epithelium (RE) over it highly infiltrated with lymphocytes. Clusters of B/plasma cells indicated by dashed lines. The crypt and the stratified squamous epithelium (SSE) on the other border of the crypt are also designated in the picture. CD1d (green), IgG/IgM (red) and nucleus (blue) staining. Arrows indicate some of the many IgG^+^/IgM^+^ B and plasma cells scattered all over the tissue and crypt. Scale bar, 104 μm. E) Representative immunofluorescence staining showing single a germinal center (GC). CD1d (green), IgG/IgM (red) and nucleus (blue) staining. Scale bar, 104 μm. F) Representative immunofluorescence staining showing a single germinal center (GC). CD1d (green), IgA (red) and nucleus (blue). Arrows indicate some of the many IgA^+^ B and plasma cells into the mesh of the reticulated epithelium (RE). Scale bar, 104 μm. Samples were examined with a Nikon Eclipse Ti-E microscope.

To visualize the lymphoid compartment, we stained for (IgM/IgG)^+^ B cells (Fig. 3). Our results confirmed previous reports (Bernstein *et al*, 2005). (IgM/IgG)^+^ B cells (and plasma cells, distinguish by their size) appearead located not only in follicles, GC an their mantle zone, but also uniformly scattered throughout the tonsillar tissues (Fig. 3A-F), generally associated in clusters when infiltrating reticular epithelial tissue (Fig. 3D). IgA^+^ B/plasma cells on the other hand, were mainly found outside the germinal centers, especially in the epithelium of crypts and rarely appear in the GCs (Fig. 3F). Also, it can be observed the protrusion of lymphocytes and plasma cells from the sites of desquamation to the lumen (Fig. 3C and D). IgG and IgM can be observed in its soluble form, accumulated on the apical surface of the epithelial layer (Fig. 3A and D). It has been previously shown that nasopharyngeal epithelium widely expresses FcRn, which mediates IgG transport across the epithelial cell layer (Heidl *et al*, 2016). On the other hand, it does not express pIgR, hence the transport of dimeric IgA is unfeasible through this kind of epithelia (Iwasaki, 2016), making IgG the predominant protective isotype in tonsils. The histological findings we described above are in agreement with those reports. Moreover, FcRn and CD1d belong to the same family of proteins, the non-classical MHC class I molecules and both have been shown to display a restriction in their expression to specific tissues, namely epithelial cells from other mucosal sites (Blumberg *et al*, 2001). Finally, follicular hyperplasia was present in all samples as well as intensively active germinal centers (Fig 4). Such hyperplasia is attributable to proliferation of BGC, as we and others have previously shown (Reis *et al.,* 2013), (Passàli *et al*, 2004), (Sarmiento Varon *et al*, 2017). In light of the results shown in previous sections, such misguided B cell response can potentially fuel the local inflammatory microenvironment, extending the impairment of immune homeostatic control.

**Figure 4.**
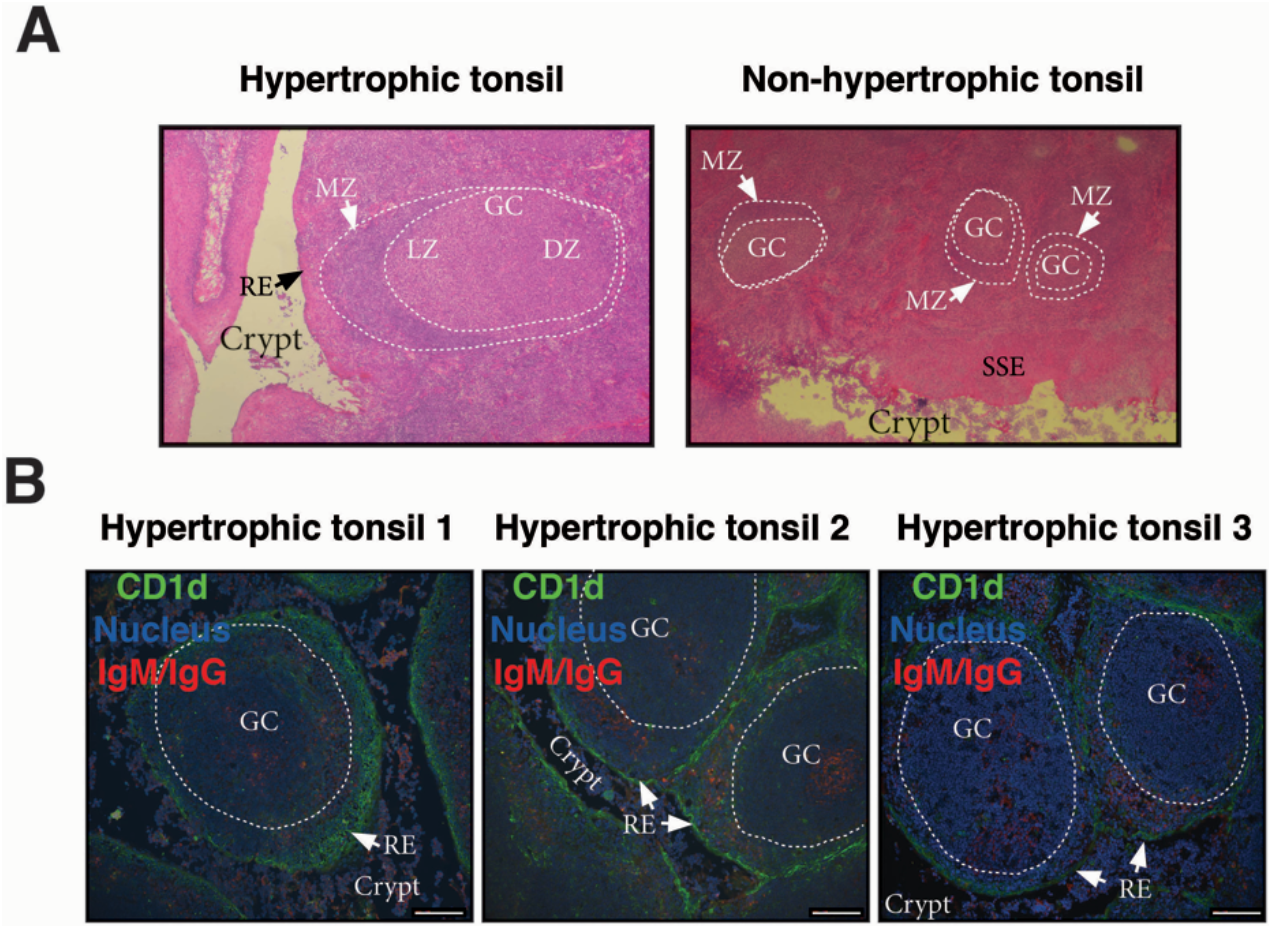
Microscopic features of germinal centers from hypertrophic tonsils. A) Comparative light microscopy of germinal centers from an hypertrophic tonsil (left) and a non-hypertrophic tonsil (right) which evidence the difference in size largely documented in literature. H.E, original magnification 40x. B) Three colour immunofluorescence staining showing germinal centers (GC) from tonsils excised from three different OSA patients (1, 2, 3, see also colour code). Note the lymphoid hyperplasia, the degree of infiltration of the reticular epithelium (RE) in all cases, and the crypts invaded by lymphoid cells (most B and plasma cells according staining). Scale bar, 104 μm. Samples were examined with a Nikon Eclipse Ti-E microscope.

### Microbiological aspects of hypertrophied tonsils

The most evident potential source of a local persistent immune response would be a local persistent infection. However, as opposed to recurrent tonsillitis, tonsillar hypertrophy has been long considered of non-infectious etiology. Actually, hypertrophied tonsils serve as non-infected control for tonsils affected by recurrent tonsillitis in a number of publications, including relevant and recent ones in which such control should be crucial (Dan *et al*, 2019). It has been generally considered that bacterial cultures from hypertrophied tonsils largely reflect oropharyngeal colonization. It is actually quite complex to distinguish infection versus colonization, provided that tonsils, being part of the mucosal immune system, are constantly exposed to the environment.

In order to try to discriminate between commensals and pathogens we started by naming the bacterial species that we could retrieve alive from the core of the samples upon cauterization of the surface, trying to identify abundant, relentless and viable populations from the clinical tissue. We cultured the core tonsillar tissue of 31 children undergoing tonsillectomy due to OSA, whose ages ranged from 2 to 15 years old. Within our cohort of patients, the frequency of tonsillectomy varied with the age, following a unimodal distribution which peaked at 6 years old (Fig 5A). All samples rendered viable bacterial cultures except for one. At phylum level, the tonsil cultures were dominated by *Firmicutes* (predominantly from the genera *Streptococcus* and *Staphylococcus* in that order), *Proteobacteria* (mostly from the genera *Neisseria* and *Haemophilus), Bacteroidetes* (principally genera *Prevotella), Actinobacteria* and *Fusobacterium* (Table 1) in agreement with recent data obtained by next generation sequence (NGS) (Fagö-Olsen *et al*, 2019).

**Figure 5.**
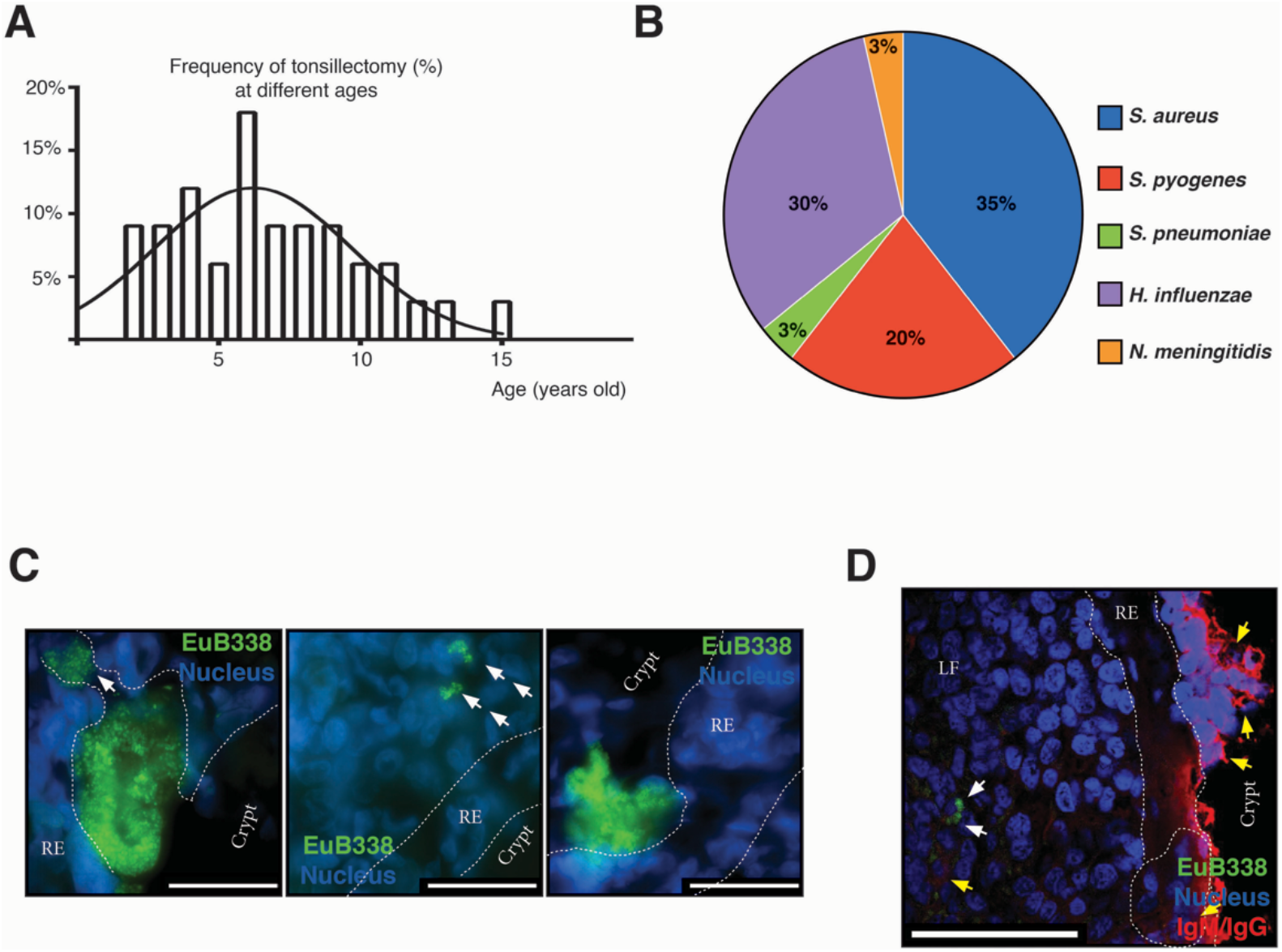
Microbiological aspects of hypertrophic tonsils. A) Age distribution of patients who had tonsillectomy due to OSA. A Gauss normal distribution fitted the observed age distribution (plotted). B) The graph shows the frequency of potential pathogenic bacterial detection in tonsils. C) Microphotographs of the bacterial aggregates in tonsillar frozen sections from different OSA patients evidenced by fluorescence *in situ* hybridization (FISH) with a general eubacterial probe (EUB338-Alexa 488, green). Host cells nuclei were stained with DAPI (blue). Dashed line represents the limits of the reticular epithelium (RE) on both margins. White arrows indicate the conglomerates of bacteria that have penetrated the epithelial barrier. Scale bar, 30 μm. Samples were examined with a Nikon Eclipse Ti-E microscope. D) Three-color immunofluorescent confocal microscopy of tonsillar frozen sections. Bacterial aggregates in tonsillar frozen sections were evidenced as in D) and are indicated with white arrows. Yellow arrows indicate B and plasma cells. Dashed line represents the limits of the reticular epithelium (RE) on both margins. Clusters of B/plasma cells are enclosed by dashed lines. LF stands for Lymphoid Follicle. Scale bar, 30 μm. Samples were examined with an Olympus FV 1000 confocal microscope.

**Table 1.**
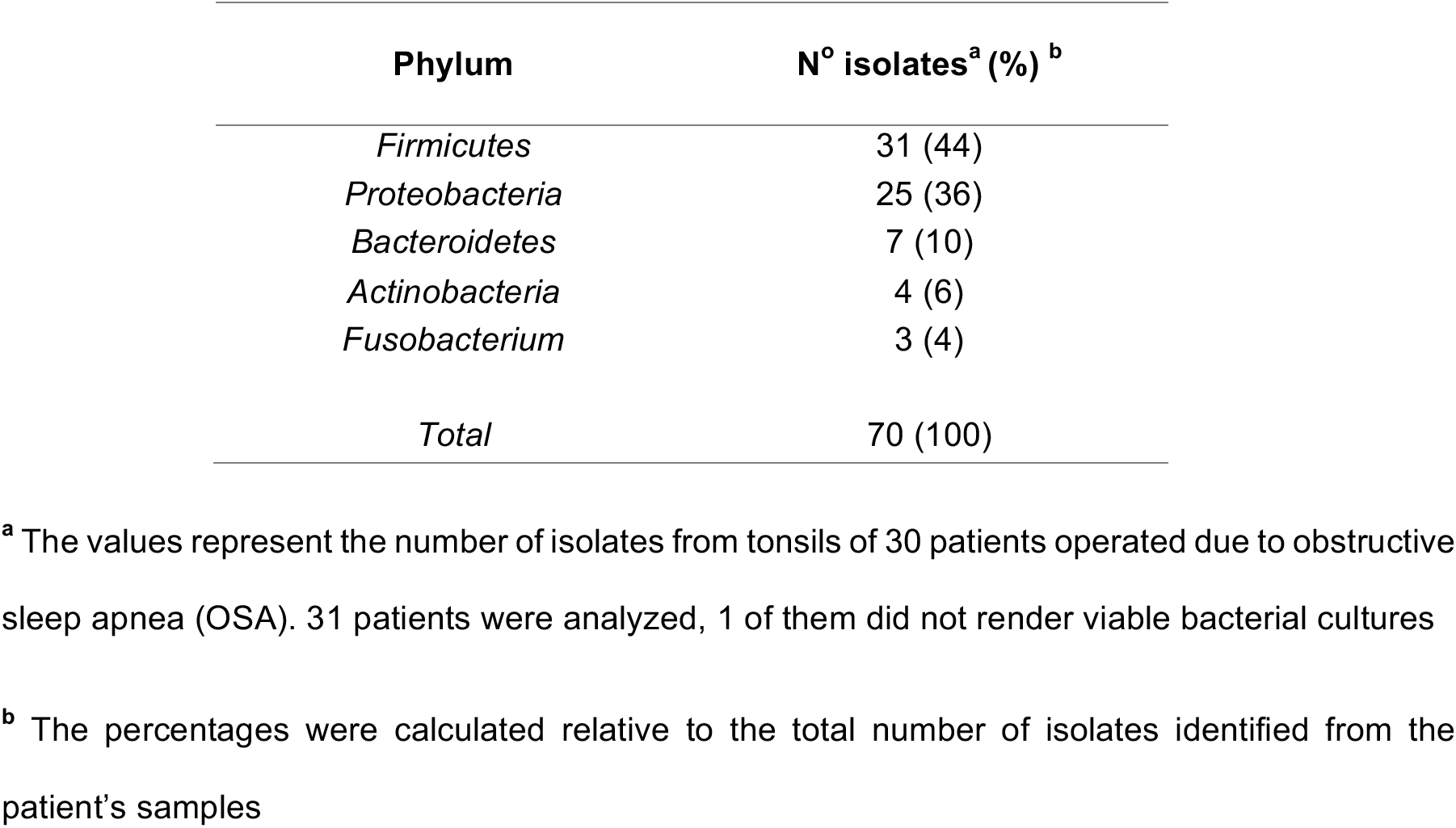
Taxonomic breakdown of isolates from the core of hypertrophied tonsils at phylum level.

| **Phylum** | **N° isolates<sup>a</sup> (%) <sup>b</sup>** |
| --- | --- |
| *Firmicutes* | 31 (44) |
| *Proteobacteria* | 25 (36) |
| *Bacteroidetes* | 7 (10) |
| *Actinobacteria* | 4 (6) |
| *Fusobacterium* | 3 (4) |
| *Total* | 70 (100) |

<sup>a</sup> The values represent the number of isolates from tonsils of 30 patients operated due to obstructive sleep apnea (OSA). 31 patients were analyzed, 1 of them did not render viable bacterial cultures
<sup>b</sup> The percentages were calculated relative to the total number of isolates identified from the patient's samples

Considering that certain microorganisms like *Streptococcus pyogenes (S. pyogenes), Streptococcus pneumoniae (S. pneumoniae), Moraxella catarrhalis (M. catarrhalis), Haemophilus influenzae (H. influenzae)* and *Staphylococcus aureus (S. aureus)* had been found either causing ear, nose and tonsil (ENT) pathology or as harmless local flora, both situations in competent hosts, we wanted to investigate their prevalence in our patients. We did not detect *M. catarrhalis* within our cohort of patients. We isolated *S. pneumoniae* from a single patient. On the other hand, *S. aureus*, isolated from over one third of patients, accounted for the most frequent potentially pathogenic bacteria among the tonsils we analysed, closely followed by *H. influenzae* (present in 30% of the patients) and *S. pyogenes* (on 20% of the samples). Bacterial isolates potentially pathogenic from the excised samples are shown in Fig 5B. In Argentina, vaccination against *H. influenzae* type b, *S. pneumoniae, Neisseria meningitidis (N. meningitidis)* and *Corynebacterium diphtheria,* which have been long known as commensal species in tonsils, is mandatory. We did not detect any patient carrying *Corynebacterium diphtheriae* but we found *Neisseria meningitidis* in the tonsils of a single patient and we did not serotype any isolate of *H. influenzae.*

In most cases we obtained co-isolates of different bacteria. Only 4 patients rendered cultures of single species, 3 of those were *Streptococcus pyogenes* group A. Co-isolates ranged from 2 to 5 species. Details of all isolated organisms from the samples are listed in Table 2. As we isolated a plethora of viable bacteria considered commensal and potential pathogenic as well, we were still unable to discriminate whether the tonsillar hypertrophy is of infectious nature.

**Table 2.** Taxonomic breakdown of isolates from the core of hypertrophied tonsils at genus and species level.

| **Aerobic bacterial isolates** | **N° patients <sup>a</sup> (%) <sup>b</sup>** |
| --- | --- |
| *Staphylococcus aureus* | 11 (35) |
| *Streptococcus viridans* | 13 (42) |
| *Streptococcus salivarius* | 2 (6) |
| *Streptococcus pneumoniae* | 1 (3) |
| *Streptococcus dysgalactiae* | 2 (6) |
| *Streptococcus pyogenes* | 6 (20) |
| *Gemella haemolysans* | 1 (3) |
| *Rothia mucilaginosa* | 3 (10) |
| *Rothia dentocariosa* | 1 (3) |
| *Haemophilus influenzae* | 9 (30) |
| *Haemophilus parainfluenzae* | 1 (3) |
| *Neisseria elongata* | 2 (6) |
| *Neisseria flavescens* | 3 (10) |
| *Neisseria subflava* | 9 (30) |
| *Neisseria meningitidis* | 1 (3) |
| *Neisseria sp* | 8 (26) |
| **Anerobic bacterial isolates** |  |
| *Solobacterium moorei* | 1 (3) |
| *Prevotella spp* | 1 (3) |
| *Prevotella oris* | 3 (10) |
| *Prevotella intermedia* | 3 (10) |
| *Prevotella nanciensis* | 2 (6) |
| *Fusobacterium necrophorum* | 1 (3) |
| *Fusobacterium nucleatum* | 1 (3) |
| *Veillonella atypica* | 1 (3) |
| *Aggregatibacter aphrophilus* | 2 (6) |
| *Corynebacterium argenteorotense* | 1 (3) |

<sup>a</sup> The values represent the number of patients that rendered the isolated designated in the column of the left
<sup>b</sup> The percentages were calculated relative to the total number of patients (31)

The basic distinction between a pathogen and a commensal is that the first one displays aggressive tools for invasion. Usually this represents that such pathogen is able to penetrate the epithelial barriers. Therefore, we set out to detect bacterial presence in tonsillar biopsies by FISH, which allowed us to investigate its distribution and organization *in situ.* We used a fluorescent universal eubacterial (EUB338) probe followed by immune-fluorescence staining on cryosections from excised tonsils. As shown in Fig. 5C and D, we detected bacterial aggregates not only associated with the surface of the epithelium but also within the lymphoid compartment, having breached the reticular epithelium. Therefore, while we cannot ascertain that the microorganisms detected *in situ* as well as through culture are the initiators of the ongoing inflammatory response, we did evidence that the chronification of the process must be certainly related to bacterial spreading beyond the normal boundaries. Also, such invasion is not associated to a single species as it can be observed in different patients harboring different colonizers, which have become pathogens, given the host’ requirement of the surgery to restore homeostasis.

## Discussion

Tonsils offer a useful model to study the interactions between the human mucosal immune system and the microbes that populates them at the epithelial level. Clinical material is not scarce, as tonsillectomy remains one of the most frequent paediatric surgeries carried out worldwide as a result of diagnosis with OSA. They rank among the secondary lymphatic organs, so every single developmental B cell subset is represented. In this study, initially we show that B cells are able to produce TNFa at similar or even higher levels than those reached by T cells, depending on the stimulating conditions. We have previously demonstrated B cell ability to rapidly produce other pro-inflammatory cytokines as IL6, IL8 and IL17 (Sarmiento Varon *et al*, 2017). Taken together, these results led us to conclude that B cell hyperplasia boost the pathological inflammatory processes and tissue alterations characteristic of OSA tonsils. Furthermore, we detected a relevant proportion of CD3^+^ cells secreting IL17, even though we did not use culture conditions capable of specifically stimulating T cells. It has been previously reported that Th17/Treg ratio measured in PBMC is positively related to the severity of OSA and serum levels of C-reactive protein (Ye *et al.,* 2012). Interestingly, there are a number of human diseases in which B cells contribute to the pathology through the acquisition of a pro-inflammatory cytokine profile and thereby supporting T cell-mediated inflammation (Ding *et al.,* 2013), (Barbosa *et al.,* 2011), (Quinn *et al.,* 2018), (DeFuria *et al*, 2013), (Zhu *et al*, 2014).

While others have reported on the expression of CD1d in different mucosal epithelial cells, (Blumberg *et al.*, 1991), (Bonish *et al.*, 2000), (Kawana *et al.*, 2008) we are the first ones to report its expression by tonsillar stratified as well as reticular epithelium. CD1d is a non-polymorphic MHC class I-like antigen-presenting molecule, restricted to lipid Ag presentation to *i*NKT cells. CD1d-binding lipid Ags from tumors as well as commensal and pathogenic microorganisms have been well characterized by many authors including ourselves (Gentilini, 2016), (**Brigl M**, 2010), (Kinjo *et al*, 2006) (De Libero *et al*, 2005). Tissue specific functions of *i*NKT cells were recently reviewed in (Crosby & Kronenberg, 2018). Of note, CD1d-mediated Ag presentation by epithelial/parenchimal cells can shape the outcome of immune responses by regulating invariant natural killer T cells (*i*NKT) function. This has been demonstrated for the case of hepatocytes, which through CD1d-lipid presentation induce *i*NKT cell tolerance in the liver, protecting from hepatic inflammation. Whereas *i*NKT cells were confirmed to be present in tonsils (Jimeno *et al*, 2019), it remains unknown their specific function at this tissue. Provided we found abundant epithelial expression of CD1d, the diversity of the tonsillar microbiota and the constant contact with environmental Ags, we suggest *i*NKT cells might be functionally as relevant at this site as they are in the gut (Brailey *et al*, 2020).

Another non-classical MHC class I molecule which has been shown to be expressed by tonsillar epithelial cells is the FcRn. Tonsillar epithelium has been classified as a type II mucosal surface (Iwasaki, 2016). In fact, such definition would fit for the stratified squamous epithelium. The specialized epithelium that covers the tonsillar crypts is only found in that location and is called either reticular or lymphoepithelium. Neither the squamous nor the reticular epithelium express pIgR, but they do express FcRn (Iwasaki, 2007). Hence, the major post switch protective Ig isotype translocated to the lumen from the tonsillar crypts is IgG. Of note, IgM and IgG are the best isotypes to activate complement. This capacity makes them particularly efficient but also more injurious to the host due to the inflammation associated with such effector immune function. Through anti (IgM/IgG) staining we evidenced the hypertrophic GCs, origin of the IgG locally produced (Fig. 4 and 5). We also stained for IgA and confirmed that IgA^+^ B cells were mainly found outside the germinal centers. Of note, it has been recently discovered in murine intestine a network of cellular interactions that drives IgM-to-IgA class switch recombination in extrafollicular B cells, compatible with the pattern we observed (Reboldi *et al*, 2016).

When studying the pathogenesis of OSA caused by hypertrophic tonsils, host immune response is only one side of the equation. The other side of it, is the microbiota populating the tonsillar mucosal surface. A diverse pool of bacteria comprising both commensal and pathogenic organisms have been isolated from OSA tonsils. In this study we worked with a cohort of patients whose ages ranged between 2 and 15 years old. In agreement with previous reports from patients elsewhere (Mattila *et al*, 2001), the frequency of tonsillectomies within that age limit peaked at 6 years old, which is coincident with the beginning of the transition from deciduous to permanent dentition for most children. Interestingly, it has been shown that such process implies the most striking change in the salivary microbiome of all ages, when considering children from birth until 18 years old (Crielaard *et al*, 2011). Choi et al. (Choi *et al*, 2020) recently demonstrated that the microbiome profiles of saliva and tonsils are largely similar both, in terms of diversity and composition, in children operated due to OSA. Given the location of tonsils at the back of the oral cavity, it was somehow expected the close relation between the two microbial groups. We could therefore speculate that the important changes in the bacterial species detected when starting to acquire permanent teeth might exert extra pressure on the tonsils of some children, being the draining immune sites first handling oral Ags. Again, probably a number of factors are responsible for the loss of homeostasis, including those of the microflora but also an altered host cell immune function. In relation to the specific bacterial species identified by culture in our samples, they are much in agreement with other studies performed either by culture or, more recently, by employing 16S rRNA gene sequencing (Table 1) (Fagö-Olsen *et al.,* 2019), (Choi *et al.,* 2020), (Viciani *et al*, 2016), (Johnston *et al*, 2018), (Jeong *et al*, 2007). In our experiments, *S. aureus* (35%), *H. influenzae* (30%) and *N. subflava* (30%) were the most prevalent bacterial species in the core of hypertrophied tonsillar tissue within our cohort of patients. They were followed by S. *Pyogenes* (20%) and different species of the genus *Streptococcus* and *Neisseria.* Finally, among the anaerobic prevalent isolates, we found species of *Prevotella, Veionella and Fusobacterium.* As we have pointed out, all these microorganisms have been isolated from around the world, from tonsils excised due to OSA and also because of recurrent tonsillitis. Clearly, there are some differences depending on the vaccination schemes operating in different countries, which might impact in differences in the prevalence within tonsils’ crypts of *S. Pneumoniae* or *N. meningitidis,* for example. However, in general, there is consensus that the bacterial species we have found are normal oropharyngeal commensals in children. A point of particular interest to discuss is whether this means that such commensal can never be injurious. Noticeable, complications in the host immune system would imply the redefinition of most commensals as pathogens. On the other hand, a change in the model of life of these bacterial communities would have the same result. In this context, we identified bacterial aggregates *in situ.* Bacteria can subsist either in planktonic or biofilm states. Planktonic cells are freely motile entities persisting in a liquid environment. In contrast, most of the bacterial species we named above (*Haemophilus, Sptaphylococcus, Streptococcus*) have long been known to have the capacity of forming biofilms. In biofilms, bacteria exist as sessile aggregates encased in a self-produced complex polysaccharide matrix attached to a surface (Chole & Faddis, 2003). The establishment of biofilms is acknowledged as a virulence factor as it allows bacteria to better resist host immune responses. While we could detect extensive aggregates attached to the surface, we did not intend to particularly identify biofilms, as we did not have samples from normal tonsils to ascertain whether biofilm presence is exclusive of pathology or not. In any case, we detect bacterial penetration through the epithelial layer to the lymphoid compartment.

To conclude, the data presented here shed light on the contribution of the B cell compartment to the inflammatory response in OSA tonsils and underscore the importance of the interplay between the host immune system and the commensal microorganisms able to switch from asymptomatic colonization to invasive disease.

## Materials and Methods

### Isolation of cells

Primary human mononuclear cells were isolated from tonsils obtained from patients (total n=40 for the different experiments) undergoing tonsillectomy due to OSA. Tonsillar mononuclear cells (TMC) were prepared as follows. Briefly, tonsils were collected in phosphate buffered saline (PBS) buffer containing 50 μg ml^-1^ amphotericin B (Richet). Tissues were chopped with a scalpel in complete medium and passed through a 70 μm-pore-size cell strainer (Falcon). TMC were purified by density gradient centrifugation with Ficoll-Hypaque (GE Healthcare). The viability of primary cells, as determined by trypan blue exclusion was greater than 99% in all preparations. Informed consent was obtained from subjects before the study. The institutional ethics committee (Clinical Hospital, School of Medicine, Buenos Aires) approved the collection and use of clinical material, conformed to the provisions of the Declaration of Helsinki (as revised in Edinburgh 2000). informed consent was obtained from all participants and/or their legal guardian/s. FACS experiments were performed with freshly isolated cells only.

### Cell culture

Primary human B cells were cultured in IMDM medium (Life Technologies) containing 10% heat-inactivated fetal calf serum, 2mM L-glutamine, 100 U/ml penicillin, 100 μg/ml streptomycin, 20 mM 4-(2-hydroxyethyl)-1-piperazineethanesulfonic acid buffer (HEPES), 1 mM sodium pyruvate and 50 μM 2-mercaptoethanol (all from Invitrogen). Human IL2 (20 ng/ml; R&D Systems) and human IL4 (20 ng/ml; R&D Systems) were added immediately before experiments also as supplements. When indicated, human recombinant hCD40L (250 ng/ml; R&D Systems) and 25 μM CpG-ODN 2006 (InvivoGen) were used. Cells were cultured at 1×10^6^ cells/ml either in 24-well culture plates (1ml) or 48-well culture plates (0.5ml).

### Antibodies and flow cytometry

Fluorochrome conjugated mAbs specific for human CD3, CD20, CD4, IL17, TNFa, IFNγ IL17 and isotype control mAbs were purchased from BD Biosciences and Biolegend. To evaluate the cytokines by intracellular staining, cells were incubated with Cytofix/Cytoperm (BD PharMingen) for 20 minutes (min) in the dark and washed with Perm/Wash solution (BD PharMingen). Following permeabilization, the cells were stained with the respective anti-cytokine mAb. Cells were acquired using FACSAria II (BD Biosciences) and analyzed with FlowJo software (Treestar). Single stained controls were used to set compensation parameters. Fluorescence minus one and isotype-matched Ab controls were used to set analysis gates. The fixable viability dye eFluor 780 was purchased from eBioscience.

### Statistical analyses

The results were analyzed using GraphPad Prism 7.0 software. The statistical analysis of the results was performed using the unpaired t test, and a p value of <0.05 was considered significant unless indicated otherwise.

### Bacterial cultures

Tonsillar samples were cultured on Columbia agar containing 5% sheep blood and chocolate agar (Laboratorios Britania, Argentina) at 37°C with 5% CO2 for 24 to 48 h. Anaerobes were culture under appropriate conditions. All isolates were identified by MALDI-TOF MS. The isolates were identified by the direct colony on plate extraction method as previously described. Mass spectra were acquired using the MALDI-TOF MS spectrometer in a linear positive mode (Microflex, Bruker Daltonics). Mass spectra were analyzed in a m/z range of 2,000 to 20,000. The MALDI Biotyper library version 3.0 and MALDI Biotyper software version 3.1 were used for bacterial identification.

### Immunofluorescence

Cryostat sections (10-20 μm thickness) of tonsils were fixed and stained with mouse anti-human CD1d (clone NOR3.2/13.17 Santa Cruz Biotechnology) followed by chicken anti-mouse IgG antibody AlexaFluor 488 (Thermo Fisher Scientific). (IgG/IgM)^+^ cells were detected by addition of goat anti-human IgM+IgG (H+L) F(ab’)2 (Jackson Immunoresearch) followed by donkey anti-goat IgG AlexaFluor 594 (Jackson Immunoresearch). IgA^+^ cells were detected by addition of goat anti-human IgA F(ab’)_2_ (InVivoGen) followed by anti-goat IgG AlexaFluor 594 (Jackson Immunoresearch. Cell nuclei were visualized with 4,6-diamidino-2-phenylindole staining (DAPI, Thermo Fisher Scientific). Finally, slides were rinsed with phosphate buffered saline, air dried, mounted in Vectashield (Vector Labs) and sealed with a glass coverslip. Samples were examined with a Nikon Eclipse Ti-E fluorescence microscope.

### Bacterial localization and immunostaining

A solution of 0.5 mg of lysozyme (Sigma)/ml in 0.1 M Tris-HCl and 0.05 M Na2EDTA was added to the cryosections, followed by incubation at 37°C for 3 h. Fixed, permeabilized tonsillar sections were then incubated in a moist chamber for 4hs at 48°C in hybridization buffer (0.9 M NaCl, 20 mM Tris-HCl [pH 7.6], 0.01% sodium dodecyl sulfate, 30% formamide) containing either the universal probe EUB388 AlexaFluor 488 labeled probe or the negative control (nonsense) NONEUB388 AlexaFluor 488. Stringent washing was performed by incubating the slide in washing buffer (20 mM Tris-HCl [pH 7.6], 0.01% sodium dodecyl sulfate, 112 mM NaCl) at 48°C for 15 min in a moist chamber a number of times, and subsequently rinsed with ddH2O for 5min. Finally, probed-hybridized tonsil cryosections were subjected to immunofluorescence staining as described above to determine the distribution of bacteria within the different tissue compartments. The hybridized slides were examined either with a FV1000 confocal microscope (Olympus) or a Nikon Eclipse Ti-E fluorescence microscope.

## Acknowledgments

We are grateful to Gastón Amable and M. Elisa Picco for their technical assistance with the immunofluorescence staining. This research was funded by the following Argentinean governmental agencies: ANPCyT (BID PICT 2015-0113) and UBA (20720170100004BA), grants to E. A. L. S. V was the recipient of a CONICET postgraduate scholarship. We are grateful to the anonymous patients and/or their parents that consent for the donation of samples for this research project.

## Author Contributions

L. S. V and J. D. R processed all the samples used in this study, performed most of the experiments and analyzed the data. R. R performed a number of experiments. L. A. B. and P. B performed cell sorting and advised on design of FACS experiments. G. B and N. S performed cryosections and H&E stainings. L.S.V, J. D. R, P. M. F and B. P critically reviewed and edited the manuscript. M. E. A and B. P provided samples. C. M. B, C. V and F. T. M. performed the microbiologic culture and identification. E.A supervised and designed research, analyzed the data and wrote the manuscript.

## References

1. Anolik J, Looney RJ, Bottaro A, Sanz I, Young F (2003) Down-regulation of CD20 on B cells upon CD40 activation. European Journal of Immunology 33: 2398–2409

2. Barbosa RR, Silva SP, Silva SL, Melo AC, Pedro E, Barbosa MP, Pereira-Santos MC, Victorino RM, Sousa AE (2011) Primary B-cell deficiencies reveal a link between human IL-17-producing CD4 T-cell homeostasis and B-cell differentiation. PLoS One 6: e22848

3. Bernstein JM, Baekkevold ES, Brandtzaeg P (2005) Chapter 90 - Immunobiology of the Tonsils and Adenoids. In: Mucosal Immunology (Third Edition), Mestecky J., Lamm M.E., McGhee J.R., Bienenstock J., Mayer L., Strober W. (eds.) pp. 1547–1572. Academic Press: Burlington

4. Blumberg RS, Terhorst C, Bleicher P, McDermott FV, Allan CH, Landau SB, Trier JS, Balk SP (1991) Expression of a nonpolymorphic MHC class I-like molecule, CD1D, by human intestinal epithelial cells. The Journal of Immunology 147: 2518–2524

5. Blumberg RS, van de Wal Y, Claypool S, Corazza N, Dickinson B, Nieuwenhuis E, Pitman R, Spiekermann G, Zhu X, Colgan S et al (2001) The multiple roles of major histocompatibility complex class-I-like molecules in mucosal immune function. Acta odontologica Scandinavica 59: 139–144

6. Bonish B, Jullien D, Dutronc Y, Huang BB, Modlin R, Spada FM, Porcelli SA, Nickoloff BJ (2000) Overexpression of CD1d by keratinocytes in psoriasis and CD1d-dependent IFN-gamma production by NK-T cells. J Immunol 165: 4076–4085

7. Boussiotis VA, Nadler LM, Strominger JL, Goldfeld AE (1994) Tumor necrosis factor alpha is an autocrine growth factor for normal human B cells. Proc Natl Acad Sci U S A 91: 7007–7011

8. Brailey PM, Lebrusant-Fernandez M, Barral P (2020) NKT cells and the regulation of intestinal immunity: a two-way street. The FEBS Journal 287: 1686–1699

9. Brandtzaeg P(2013) Secretory immunity with special reference to the oral cavity. Journal of oral microbiology 5

10. Brigl M BM (2010) How invariant natural killer T cells respond to infection by recognizingmicrobial or endogenous lipid antigens. Semin Immunol 22: 79–86

11. Choi DH, Park J, Choi JK, Lee KE, Lee WH, Yang J, Lee JY, Park YJ, Oh C, Won H-R et al (2020) Association between the microbiomes of tonsil and saliva samples isolated from pediatric patients subjected to tonsillectomy for the treatment of tonsillar hyperplasia. Experimental & Molecular Medicine

12. Chole RA, Faddis BT (2003) Anatomical evidence of microbial biofilms in tonsillar tissues: a possible mechanism to explain chronicity. Arch Otolaryngol Head Neck Surg 129: 634–636

13. Crielaard W, Zaura E, Schuller AA, Huse SM, Montijn RC, Keijser BJ (2011) Exploring the oral microbiota of children at various developmental stages of their dentition in the relation to their oral health. BMC medical genomics 4: 22

14. Crosby CM, Kronenberg M (2018) Tissue-specific functions of invariant natural killer T cells. Nat Rev Immunol 18: 559–574

15. Dan JM, Havenar-Daughton C, Kendric K, Al-Kolla R, Kaushik K, Rosales SL, Anderson EL, LaRock CN, Vijayanand P, Seumois G et al (2019) Recurrent group A Streptococcus tonsillitis is an immunosusceptibility disease involving antibody deficiency and aberrant T(FH) cells. Sci Transl Med 11

16. De Libero G, Moran AP, Gober HJ, Rossy E, Shamshiev A, Chelnokova O, Mazorra Z, Vendetti S, Sacchi A, Prendergast MM et al (2005) Bacterial infections promote T cell recognition of self-glycolipids. Immunity 22: 763–772

17. DeFuria J, Belkina AC, Jagannathan-Bogdan M, Snyder-Cappione J, Carr JD, Nersesova YR, Markham D, Strissel KJ, Watkins AA, Zhu M et al (2013) B cells promote inflammation in obesity and type 2 diabetes through regulation of T-cell function and an inflammatory cytokine profile. Proc Natl Acad Sci U S A 110: 5133–5138

18. Ding Y, Li J, Wu Q, Yang P, Luo B, Xie S, Druey KM, Zajac AJ, Hsu H-C, Mountz JD (2013) IL-17RA Is Essential for Optimal Localization of Follicular Th Cells in the Germinal Center Light Zone To Promote Autoantibody-Producing B Cells. The Journal of Immunology 191: 1614

19. Endres R, Alimzhanov MB, Plitz T, Fütterer A, Kosco-Vilbois MH, Nedospasov SA, Rajewsky K, Pfeffer K (1999) Mature Follicular Dendritic Cell Networks Depend on Expression of Lymphotoxin β Receptor by Radioresistant Stromal Cells and of Lymphotoxin β and Tumor Necrosis Factor by B Cells. Journal of Experimental Medicine 189: 159–168

20. Fagö-Olsen H, Dines LM, Sørensen CH, Jensen A (2019) The Adenoids but Not the Palatine Tonsils Serve as a Reservoir for Bacteria Associated with Secretory Otitis Media in Small Children. mSystems 4: e00169–00118

21. Fillatreau S (2018) B cells and their cytokine activities implications in human diseases. Clinical immunology (Orlando, Fla) 186: 26–31

22. Fu Y-X, Chaplin DD (1999) DEVELOPMENT AND MATURATION OF SECONDARY LYMPHOID TISSUES. Annual Review of Immunology 17: 399–433

23. Gentilini MP, ME; Fernández, PM; Fainboim, L and Arana, E (2016) The tumor antigen N-Glycolyl-GM3 is a human CD1d ligand capable of mediating B cell and natural killer T cell interaction Cancer Immunology, immunotherapy 65: 551–562

24. Gonzalez M, Mackay F, Browning JL, Kosco-Vilbois MH, Noelle RJ (1998) The Sequential Role of Lymphotoxin and B Cells in the Development of Splenic Follicles. Journal of Experimental Medicine 187: 997–1007

25. Heidl S, Ellinger I, Niederberger V, Waltl EE, Fuchs R (2016) Localization of the human neonatal Fc receptor (FcRn) in human nasal epithelium. Protoplasma 253: 1557–1564

26. Huang YS, Guilleminault C, Hwang FM, Cheng C, Lin CH, Li HY, Lee LA (2016) Inflammatory cytokines in pediatric obstructive sleep apnea. Medicine (Baltimore) 95: e4944

27. Iwasaki A (2007) Mucosal dendritic cells. Annu Rev Immunol 25: 381–418

28. Iwasaki A (2016) Exploiting Mucosal Immunity for Antiviral Vaccines. Annu Rev Immunol 34: 575–608

29. Jeong JH, Lee DW, Ryu RA, Lee YS, Lee SH, Kang JO, Tae K (2007) Bacteriologic comparison of tonsil core in recurrent tonsillitis and tonsillar hypertrophy. Laryngoscope 117: 2146–2151

30. Jimeno R, Lebrusant-Fernandez M, Margreitter C, Lucas B, Veerapen N, Kelly G, Besra GS, Fraternali F, Spencer J, Anderson G et al (2019) Tissue-specific shaping of the TCR repertoire and antigen specificity of iNKT cells. Elife 8

31. Johnston J, Hoggard M, Biswas K, Astudillo-García C, Waldvogel-Thurlow S, Radcliff FJ, Mahadevan M, Douglas RG (2018) The bacterial community and local lymphocyte response are markedly different in patients with recurrent tonsillitis compared to obstructive sleep apnoea. Int J Pediatr Otorhinolaryngol 113: 281–288

32. Kawana K, Matsumoto J, Miura S, Shen L, Kawana Y, Nagamatsu T, Yasugi T, Fujii T, Yang H, Quayle AJ et al (2008) Expression of CD1d and ligand-induced cytokine production are tissue specific in mucosal epithelia of the human lower reproductive tract. Infection and immunity 76: 3011–3018

33. Kheirandish-Gozal L, Gozal D (2019) Obstructive Sleep Apnea and Inflammation: Proof of Concept Based on Two Illustrative Cytokines. International journal of molecular sciences 20

34. Kinjo Y, Tupin E, Wu D, Fujio M, Garcia-Navarro R, Benhnia MR, Zajonc DM, Ben-Menachem G, Ainge GD, Painter GF et al (2006) Natural killer T cells recognize diacylglycerol antigens from pathogenic bacteria. Nat Immunol 7: 978–986

35. Mattila PS, Tahkokallio O, Tarkkanen J, Pitkäniemi J, Karvonen M, Tuomilehto J (2001) Causes of tonsillar disease and frequency of tonsillectomy operations. Arch Otolaryngol Head Neck Surg 127: 37–44

36. Olah I, Glick B (1978) The number and size of the follicular epithelium (FE) and follicles in the bursa of Fabricius. Poultry science 57: 1445–1450

37. Parekh VV, Lalani S, Kim S, Halder R, Azuma M, Yagita H, Kumar V, Wu L, Kaer LV (2009) PD-1/PD-L blockade prevents anergy induction and enhances the anti-tumor activities of glycolipid-activated invariant NKT cells. J Immunol 182: 2816–2826

38. Passàli D, Damiani V, Passàli GC, Passàli FM, Boccazzi A, Bellussi L (2004) Structural and immunological characteristics of chronically inflamed adenotonsillar tissue in childhood. Clinical and diagnostic laboratory immunology 11: 1154–1157

39. Perez ME, Billordo LA, Baz P, Fainboim L, Arana E (2014) Human memory B cells isolated from blood and tonsils are functionally distinctive. Immunol Cell Biol 92: 882–887

40. Quinn JL, Kumar G, Agasing A, Ko RM, Axtell RC (2018) Role of TFH Cells in Promoting T Helper 17-Induced Neuroinflammation. Front Immunol 9: 382

41. Reboldi A, Arnon TI, Rodda LB, Atakilit A, Sheppard D, Cyster JG (2016) IgA production requires B cell interaction with subepithelial dendritic cells in Peyer’s patches. Science 352: aaf4822

42. Reis LG, Almeida EC, da Silva JC, Pereira Gde A, Barbosa Vde F, Etchebehere RM (2013) Tonsillar hyperplasia and recurrent tonsillitis: clinical-histological correlation. Braz J Otorhinolaryngol 79: 603–608

43. Sarmiento Varon L; De Rosa, J; Machicote, A; Billordo, LA; Baz, P; Fernández, PM; Kaimen Maciel, I; Blanco, A; Arana E (2017) Characterization of tonsillar IL10 secreting B cells and their role in the pathophysiology of tonsillar hypertrophy. Scientific reports 7, 1077

44. Valentine MA, Cotner T, Gaur L, Torres R, Clark EA (1987) Expression of the human B-cell surface protein CD20: alteration by phorbol 12-myristate 13-acetate. Proceedings of the National Academy of Sciences 84: 8085–8089

45. Viciani E, Montagnani F, Tavarini S, Tordini G, Maccari S, Morandi M, Faenzi E, Biagini C, Romano A, Salerni L et al (2016) Paediatric obstructive sleep apnoea syndrome (OSAS) is associated with tonsil colonisation by Streptococcus pyogenes. Sci Rep 6: 20609

46. Yamashita K, Ichimiya S, Kamekura R, Nagaya T, Jitsukawa S, Matsumiya H, Takano K, Himi T (2016) Studies of Tonsils in Basic and Clinical Perspectives: From the Past to the Future. Adv Otorhinolaryngol 77: 119–124

47. Ye J, Liu H, Zhang G, Li P, Wang Z, Huang S, Yang Q, Li Y (2012) The Treg/Th17 Imbalance in Patients with Obstructive Sleep Apnoea Syndrome. Mediators of Inflammation 2012: 815308

48. Zhu M, Belkina AC, DeFuria J, Carr JD, Van Dyke TE, Gyurko R, Nikolajczyk BS (2014) B cells promote obesity-associated periodontitis and oral pathogen-associated inflammation. J Leukoc Biol 96: 349–357

